# Pf-PeptideFilter: An Interactive Catalogue of Peptide Vaccine Candidates for *Plasmodium falciparum*

**DOI:** 10.64898/2025.12.15.694343

**Authors:** Andrew J. Balmer, Chiyun Lee, Richard D. Pearson, Cristina V. Ariani, Jacob Almagro Garcia

**Affiliations:** Wellcome Sanger Institute, Wellcome Genome Campus, Hinxton, Cambridge, CB10 1SA, UK; Liverpool School of Tropical Medicine, Pembroke Place, Liverpool, L3 5QA, UK

## Abstract

In the last decade, there has been substantial progress in vaccine development for malaria, with two WHO-recommended vaccines now available–RTS,S/AS01 (Mosquirix) and R21/Matrix-M. While these vaccines represent significant milestones, their partial efficacy underscores the need to develop next-generation vaccine approaches capable of broader, longer-lasting protection. Peptide vaccines are a promising strategy, as they focus on short, synthetic peptides specifically selected to trigger a T-cell immune response. However, developing effective peptide vaccines for malaria remains challenging, due to the extensive genetic diversity of *Plasmodium falciparum.* Currently, tools for comparing and prioritising immunogenic targets remain limited, and existing approaches overlook key factors, including genetic variation, lifecycle-stage expression, and peptide-level properties, each of which are critical for vaccine efficacy. To address these gaps, we developed Pf-PeptideFilter, an interactive web-based application and curated database designed for filtering peptide vaccine targets across the *P. falciparum* genome. By integrating population-scale genomic information (Pf7), liver-stage transcriptomics, and other biological annotations, Pf-PeptideFilter enables users to interactively filter potential vaccine candidates from the *P. falciparum* genome, based on several criteria relevant to vaccine design. The app allows users to prioritise candidate genes and peptides with immunogenic potential and export shortlists for incorporation into experimental workflows, offering an interactive platform for rational peptide vaccine design in *P. falciparum*.

**Author Summary:** Malaria vaccines are challenging to design because the parasite responsible for most infections, *Plasmodium falciparum*, is highly genetically diverse. If a vaccine targets part of the parasite proteome that varies between strains, it may protect against only a subset of parasites and perform poorly across regions. Peptide vaccines offer a promising approach, by focusing immune responses on short, synthetic peptides selected to trigger T-cell immunity. However, identifying which peptides to include in a vaccine is difficult, as the parasite genome contains thousands of genes and huge numbers of potential peptide targets. Here, we introduce Pf-PeptideFilter, an interactive tool that helps researchers prioritise peptide vaccine candidates using a series of biologically relevant filters. We integrate population-scale genome variation from thousands of parasite samples with information on gene expression during liver infection, similarity to human proteins, and conservation across related parasite species. Users can adjust filtering thresholds and immediately see how these choices affect which genes and peptides are retained. Pf-PeptideFilter generates downloadable shortlists designed to support experimental follow-up, providing a robust and transparent platform for the early stages of vaccine design.

## Introduction

Malaria remains a significant global health challenge, causing an estimated 263 million cases and 597,000 deaths annually worldwide (1,2). Among human malaria parasites, *Plasmodium falciparum* is the most common, and is responsible for the greatest mortality burden (3). Despite initial progress in mitigating its spread, reductions in the overall burden of the parasite have plateaued over the past decade, highlighting an urgent need for innovative control strategies (4). Since 2022, traditional antimalarial strategies (e.g. antimalarials, residual spraying, and insecticide-treated bednets) have been complemented by vaccines, which have now emerged as critical tools in malaria control (4–6). As of 2024, two primary vaccines are in use; the RTS,S/AS01 (Mosquirix) and R21/Matrix-M (6,7). The RTS,S vaccine provides partial protection, with an efficacy of approximately 30-40% in young children (8,9), while the newer R21 vaccine has demonstrated higher efficacy, with around 78% protection in the same demographic (6,10,11). While these results are promising, there are several caveats–the protection offered by these vaccines can wane over time, and varies by both age and vaccine schedule (7,12,13).

Although the success of these vaccines represent significant milestones in reducing the global malaria burden, they also underscore the need for the next generation of vaccine solutions that can offer longer-lasting and broader protection (14,15). Peptide vaccines represent a promising approach. These vaccines differ from existing malaria vaccines such as RTS,S and R21, which use recombinant proteins or virus-like particles, as they instead use short, synthetic peptides, specifically selected to induce targeted T-cell immune responses. Peptide vaccines offer several advantages over conventional approaches, as they are easier and cheaper to manufacture, highly specific, and can be customised to match immune profiles in specific populations (16,17). While these approaches have not been widely used for *P. falciparum*, they have shown strong potential in trials with other pathogens, such as hepatitis C virus and SARS-CoV-2 (18,19). These results show that peptide-based vaccine design could strongly influence real-world vaccine pipelines, and with systematic candidate selection–making use of the extensive genomic surveillance data available for *P. falciparum*–it could play a major role for malaria as well. For instance, peptide vaccines may work well as part of a multi-component vaccine for malaria, working in synergy with B-cell immunity to reduce the likelihood of immune evasion, enhancing overall vaccine efficacy and providing tailored protection across diverse parasite populations (20–22). Together, these features could make them a highly adaptable strategy for mitigating malaria burden in endemic countries (23–25).

However, identifying effective peptide vaccine candidates for malaria is challenging. This is because *P. falciparum* is highly genetically and antigenically diverse, and its genome encompasses almost 5,000 genes. Not all peptides within its proteome are presented effectively by the immune system, and its genetic diversity means some peptides may only be present among a subset of parasite populations. This poses a significant challenge for vaccine development, as including variable or non-immunologic peptides in a vaccine would compromise its efficacy, by only targeting a subset of parasites (26–28). Similarly, geographic variability among parasite populations adds further complexity, as peptides effective in one region may perform poorly in another (29). Expression of genes can also vary across the parasite’s life-cycle, meaning targets need to be relevant to critical infection stages, such as the pre-erythrocytic liver stage (27,30,31). Lastly, homology with human proteins further reduces the pool of viable targets, due to concerns about cross-reactivity (32,33). Effective vaccine candidates must therefore meet several criteria simultaneously; they need to target conserved genomic regions, be relevant to high-burden populations, be expressed in critical infection stages, and elicit robust immune responses with minimal homology to human proteins.

Addressing all of these challenges requires a detailed catalogue of potential vaccine targets. However, the size of the *P. falciparum* genome makes comprehensive experimental screening impractical. We therefore need computational tools that can systematically filter and rank candidate peptides using genomic and proteomic data. These shortlists can then be validated experimentally to assess their suitability as vaccine candidates, using closely related model *Plasmodium* species (34,35). Although large-scale databases such as MalariaGEN Pf7, PlasmoDB, and gene expression datasets are invaluable, these tools currently lack the ability to integrate and filter datasets using multiple criteria (29–31). Similarly, while several computational tools have been developed to prioritise vaccine candidates in other organisms, these methods have not been applied to *P. falciparum*, do not make use of the extensive data now publicly available, and do not systematically filter all potential proteomic candidates using the multiple factors required (32,36–40). Developing these tools, and making them capable of filtering candidates across the entire *P. falciparum* proteome, would enable researchers to maximise the benefit of the large-scale genomic resources currently available, allowing the generation of ranked catalogues of potential vaccine targets (25,36,41).

To address these gaps, we developed Pf-PeptideFilter–a database and customisable tool for identifying potential vaccine candidates in *P. falciparum*. By leveraging the Pf7 dataset–encompassing over 16,000 *Plasmodium* genomes–one of the largest available non-human eukaryotic genomic resources, Pf-PeptideFilter offers systematic prioritisation of peptide vaccine candidates with unprecedented resolution. Pf-PeptideFilter enables users to filter genes and peptides using a range of key metrics, including; sequence conservation, gene expression profiles, cross-species homology, and similarity to human proteins, providing a generalisable tool for genomic exploration. Together, these filters provide the user with an adaptable catalogue of potential vaccine candidates, offering the ability to download and incorporate customisable output datasets into experimental workflows. By integrating Pf-PeptideFilter into an easy to use Streamlit application (42), we demonstrate its flexibility and functionality through a series of example analyses. With its open-access design, Pf-PeptideFilter has the potential to significantly advance rational vaccine development, making it an invaluable contribution to global malaria control efforts. Pf-PeptideFilter is free to use, and publicly accessible at https://pf-peptidefilter.streamlit.app/.

## Methods

### Data Sources and Pre-processing

Pf-PeptideFilter uses an aggregated database of genomic, transcriptomic, and immunological datasets to systematically filter vaccine candidates in *Plasmodium falciparum*. The primary dataset upon which this database is built is Pf7, as aggregated by MalariaGEN network, which includes high-quality whole-genome sequences from 16,203 *P. falciparum* isolates (29). This dataset provides a detailed catalogue of parasite genetic diversity, allowing precise evaluation of global sequence variation. To create a filter which prioritises genes expressed during infection, we also incorporated liver-stage transcriptomic data from Zanghi et al. (2025), which quantified parasite gene expression in hepatocytes (43).

To generate an initial set of peptide candidates across the *P. falciparum* reference genome (44), we used a sliding-window approach–extracting overlapping 20-amino-acid sequences with a 10-amino-acid offset. In total, this led to a total of 379,325 peptides within 4,937 genes. These peptide sizes were chosen to balance coverage of both MHC Class I (8-11 residues) and Class II (12-25 residues) epitopes, while maintaining computational efficiency and flexibility during downstream validation. Notably, although liver-stage peptide vaccines primarily rely on Class-I epitopes, structuring the peptide library in this way ensures compatibility with future Class-II analyses, and avoids the need for separate peptide libraries for each MHC class.

### Filtering Criteria

Pf-PeptideFilter refines the initial candidate selection through a sequential filtering process, excluding non-viable targets based on a series of filters (Table 1). Metrics are computed at the gene or peptide levels, depending on the criterion (see Supplementary Table 1). For example, gene expression is evaluated at the gene level, whereas human identity is assessed per peptide and averaged across genes. Strain conservation can be assessed at both gene and peptide levels. At the peptide level, it is calculated using the minimum frequency of the amino acid haplotype for each peptide among all Pf7 African samples. This is then extrapolated to each gene using the percentage of peptides in a gene which pass the given filter, or alternatively, the full gene haplotype. Pf-PeptideFilter provides filtering criteria for each metric to allow users to customise filter thresholds depending on their own research priorities.

**Table 1.** Brief description of the filtering stages in Pf-PeptideFilter, for more detail see Supplementary Table 1.

| Stage | Criterion | Description |
| --- | --- | --- |
| **Filtering** | Human Identity | To reduce the risk of autoimmunity, Pf-PeptideFilter can exclude peptides with high similarity to human exons. Peptide similarity is estimated using sequence identity to human peptides calculated using BLAST against human protein-coding regions (45). Filtering is performed at the peptide level and summarised per gene. |
|  | Strain Conservation | Vaccine targets with high sequence variability can lead to immune escape through genetic variation. Conservation was therefore assessed using the Pf7 dataset (16,203 high-quality genomes), by calculating peptide haplotype frequency across 8492 African samples (29). Genes and peptides with excess diversity can be excluded using percentile-based thresholds. |
|  | Indel Frequency | Similar to strain conservation, genes containing sequence variability in the form of insertions or deletion mutations (indels) are more likely to lead to immune escape. Moreover, indels can produce unstable peptides, limiting their usefulness as vaccine candidates. Pf-PeptideFilter can remove genes containing high-frequency indels, including those causing frameshifts. Indel frequencies were computed using the Pf7 dataset, allowing genes to be excluded using adjustable, percentile-based thresholds (29). |
|  | Gene Expression | To ensure candidates are expressed during infection, genes with low liver-stage expression can be removed. RNA-seq data from Zanghi et al. (2025) were used to identify expressed genes across days 2, 4, 5 and 6 in hepatocyte samples (43). Pf-PeptideFilter allows users to recall gene expression using adjustable criteria—by specifying the minimum counts-per-million (CPM) threshold and the number of biological replicates that must meet this cutoff (e.g., at least one replicate, at least two, or all replicates). This enables users to adopt more or less conservative expression-calling rules |
|  |  | depending on their research needs. |
|  | Cross-Species Homology | To prioritise targets suitable for cross-species validation in animal models, genes lacking one-to-one orthologs in *P. vivax*, *P. berghei*, or *P. knowlesi* can be excluded. Homology was determined using pre-computed reciprocal BLAST searches (30). |

We chose these filtering criteria to balance biological relevance with practical considerations for vaccine development. We therefore incorporated several metrics, including human identity, strain conservation, indel frequency, gene expression and gene homology, a full explanation of which can be found in Figure 1 and Table 1. Briefly, to reduce the risk of autoimmune cross-reactivity, sequences were compared to human exons using BLAST-based alignment, allowing filtering of peptides with high sequence identity to human exons. To ensure broad applicability across malaria endemic populations, we assessed strain conservation by calculating peptide haplotype frequencies across 8492 African isolates from the Pf7 dataset (29). Indel frequencies were also estimated using Pf7 data, allowing exclusion of genes disrupted by high-frequency insertions or deletions. We also included gene expression filtering to subset candidates based on expression profile during liver-stage infection, the phase in which parasites are more likely to trigger T-cell immunity. Finally, to identify genes suitable for cross-species validation *in vitro*, we included a filter to exclude genes lacking one-to-one orthologs in *P. vivax*, *P. berghei*, or *P. knowlesi,* using reciprocal BLAST comparisons from PlasmoDB to make these determinations (30).

**Figure 1.**
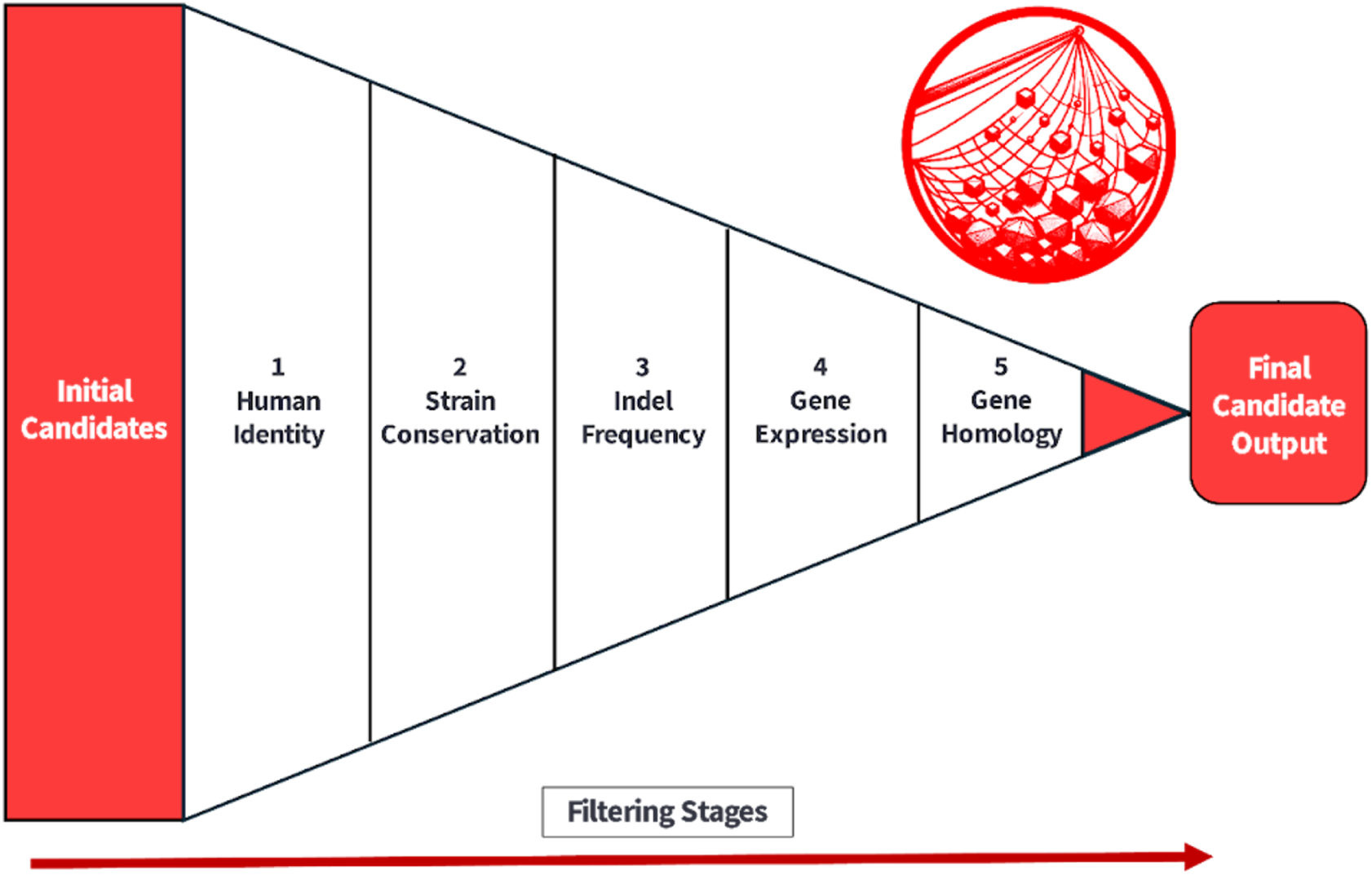
Schematic representation of the Pf-PeptideFilter filtering pipeline. The initial list of candidate genes and peptides undergoes filtering using five key criteria: 1) human identity, 2) strain conservation, 3) indel frequency, 4) gene expression, and 5) gene homology. The final candidate output includes all genes which passed the filtering criteria, and is exportable as a compressed Excel or CSV file.

### Implementation and Computational Workflow

As with previous genomic exploration tools from MalariaGEN, such as Pf-Haploatlas, Pf-PeptideFilter is implemented as an interactive web application using the Streamlit framework, with backend processing and data integration written in Python (42,46,47). To optimise performance and minimise computational burden for users, all primary metrics, including strain conservation, and human homology, are pre-computed and stored in a structured database. The app interface therefore allows dynamic adjustment of filtering thresholds utilising this database, providing the user with real-time updates and visualisation of candidate properties. In addition to candidate filtering, Pf-PeptideFilter provides an ‘enrichment analysis’ of cellular component classes for the retained gene set, based on curated Gene Ontology/PlasmoDB annotations. This analysis allows users to qualitatively assess whether filtering choices have introduced localisation biases among filtered genes (e.g. enrichment for membrane or exported proteins).

Via the app, users can preview filtered results, inspect gene-peptide filtering, and export candidate lists in CSV or Excel formats. Following filtering, two files are generated: one at the gene level, containing reference and dominant haplotypes alongside comma-separated peptide lists; and one at the peptide level, with one row per peptide, including both reference and dominant sequences. All peptides from each gene are retained in the output, regardless of the individual filtering result for a given peptide. Therefore, rather than simply excluding peptides in the output, each peptide is instead given a classifier highlighting their exclusion status, as this ensures easier compatibility with downstream comparative or immunological workflows. For large datasets (e.g. >20,000 peptides), compressed download formats are available to facilitate efficient data handling. Pf-PeptideFilter is publicly accessible at https://pf-peptidefilter.streamlit.app/.

## Results

To demonstrate Pf-PeptideFilter’s utility and flexibility, we evaluated how different filtering criteria affected the retention of candidate genes and peptides. As a first step, we applied a single, conservation-based filter and assessed its impact on gene retention. As mentioned above, amino acid haplotype frequencies were computed for each peptide using data for 8492 African isolates from the Pf7 dataset (29). When configuring Pf-PeptideFilter to retain only those genes for which at least 95% of peptides exhibited a dominant haplotype (i.e. a haplotype present in 95% of isolates or more), a total of 2,107 genes and 84,556 peptides were retained (of an initial 4,937 genes and 379,325 peptides) (Figure 2). Increasing the stringency of the filter–by requiring a greater proportion of peptides per gene to meet the threshold (99%)–resulted in a further reduction of retained genes, with 1,517 (42,208 peptides) meeting these criteria (a decrease of 28%). This relationship followed a smooth decay curve with increasing stringency (Supplementary Figure S1), illustrating the trade-off between filtering stringency and candidate pool size.

**Figure 2.**
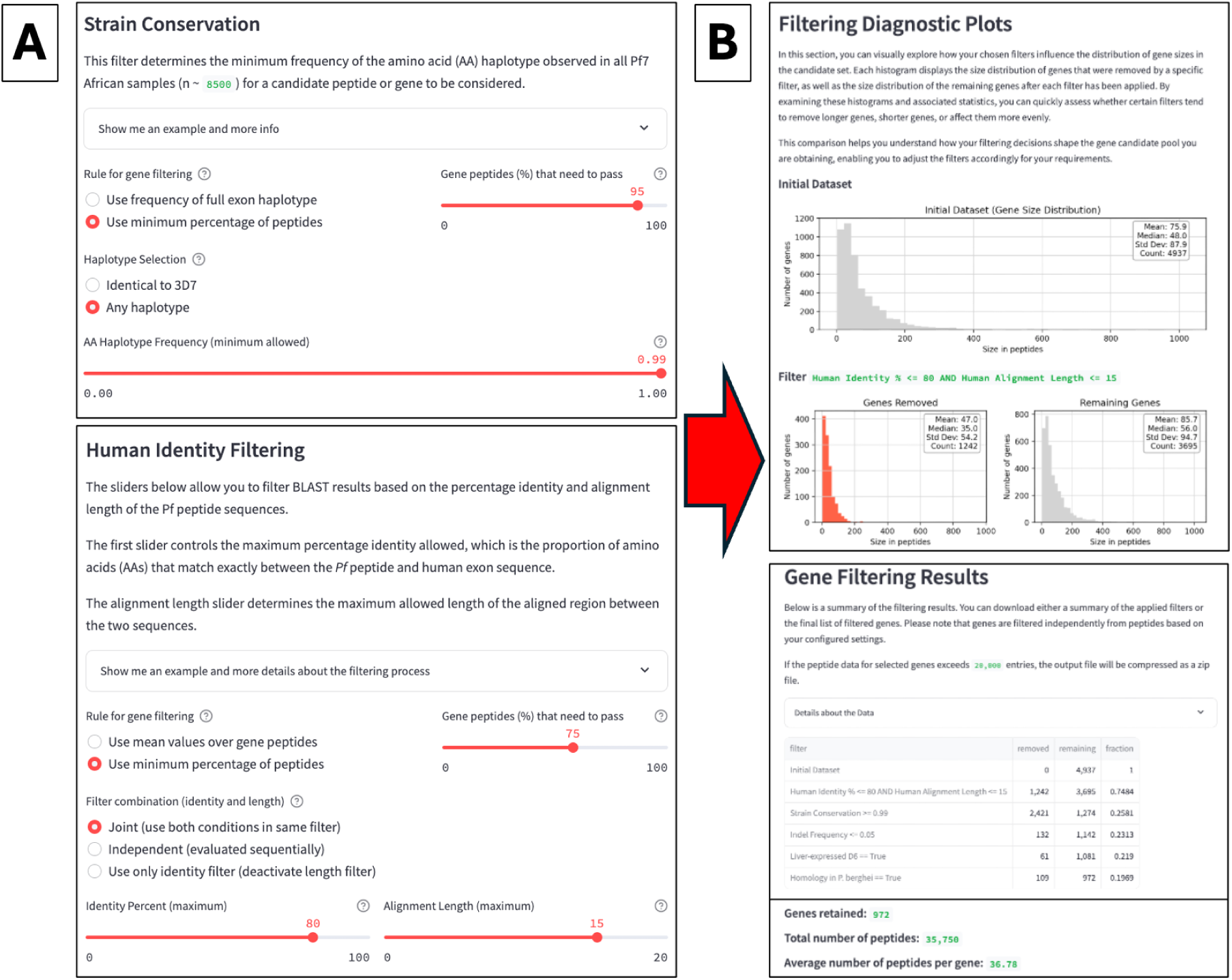
A) Pf-PeptideFilter provides a customisable user interface with which to apply sequential filtering metrics to the *P. falciparum* genome, offering tailored peptide vaccine candidate libraries for downstream experimental or computational validation. Users define the percentage of a gene’s peptides that must pass the criterion to retain that gene, between 50-100% (default: 95%), allowing researchers to calibrate stringency depending on their requirements. Filtering parameters can be modified dynamically, allowing the application of flexible, percentile-based thresholds (Figure 2). For each numeric criterion, users can define the minimum proportion of peptides that must satisfy the filter condition for the corresponding gene to be retained. B) Once filtering has been applied, the app outputs a series of diagnostic plots (see Figure 3 for more details), as well as the final results in both gene and peptide format, where the results of the sequential filtering can be observed. Candidate sets can then be downloaded as an Excel or CSV file.

We then evaluated the impact of combining multiple filters. Applying both the strain conservation and human identity filters simultaneously–i.e. by requiring 95% of peptides per gene to exceed a 99% conservation threshold and remain below 80% identity to any human exon with a maximum alignment length of 15 amino acids–reduced the number of retained genes to 1,274 (Figure 2). As expected, adding filters for indel frequency, liver-stage gene expression, and cross-species homology (presence of one-to-one orthologs in *P. vivax*, and *P. berghei*) further narrows this candidate set. When all five filters were applied using default thresholds (human identity <= 80%, strain conservation >= 99%, indel frequency < 5%. homology with *P. vivax*, gene expression in day 4 of liver stage infection), the final filtered dataset comprised 253 genes and 10,088 peptides–representing a shortlist of potential vaccine targets suitable for experimental validation.

Among the filtered genes were several whose products have been studied previously as potential components of multicomponent malaria vaccines–including SPATR (a sporozoite surface protein involved in hepatocyte invasion), TCTP (a translationally controlled tumor protein previously shown to elicit immune responses), and profilin (an actin-binding protein essential for parasite motility and invasion) (see Discussion). Importantly, these examples are given merely to illustrate how Pf-PeptideFilter can recover known vaccine-related antigens, rather than as endorsements of particular candidates. The tool also shortlisted a number of candidates from genes not previously associated with vaccine development, such as GAPM2 (a glideosome-associated protein), TOM40 (a mitochondrial import channel), USP13 (a ubiquitin-specific protease) and HSP110c (a heat-shock protein). These candidates are highlighted as cases where Pf-PeptideFilter filters genes with clear functional roles and low population variability, but would require further experimental validation before consideration as vaccine targets.

We designed Pf-PeptideFilter’s thresholding system to allow users to interactively calibrate the trade-off between candidate inclusivity and selection rigor. Filtering thresholds can be dynamically adjusted to alter the proportion of peptides within each gene required to pass a given criterion, allowing users to customise stringency (e.g. by setting a threshold from 50% to 100% of peptides per gene) (Figure 2). Real-time feedback allows users to monitor how each criterion affects the retention of candidate peptides and genes, enabling more informed filter calibration. To support user interpretation of biases, Pf-PeptideFilter provides diagnostic plots comparing length distributions between retained and excluded sets under each filter setting, as well as gene-enrichment information on the filtered genes. These metrics allow users to assess and adjust filtering parameters in a more informed and interpretable manner (Figure 3). For example, under our strain conservation filtering scenario, we observed that longer genes were more likely to be excluded at higher stringency thresholds, owing to their greater likelihood of containing variable regions (Figure 3).

**Figure 3.**
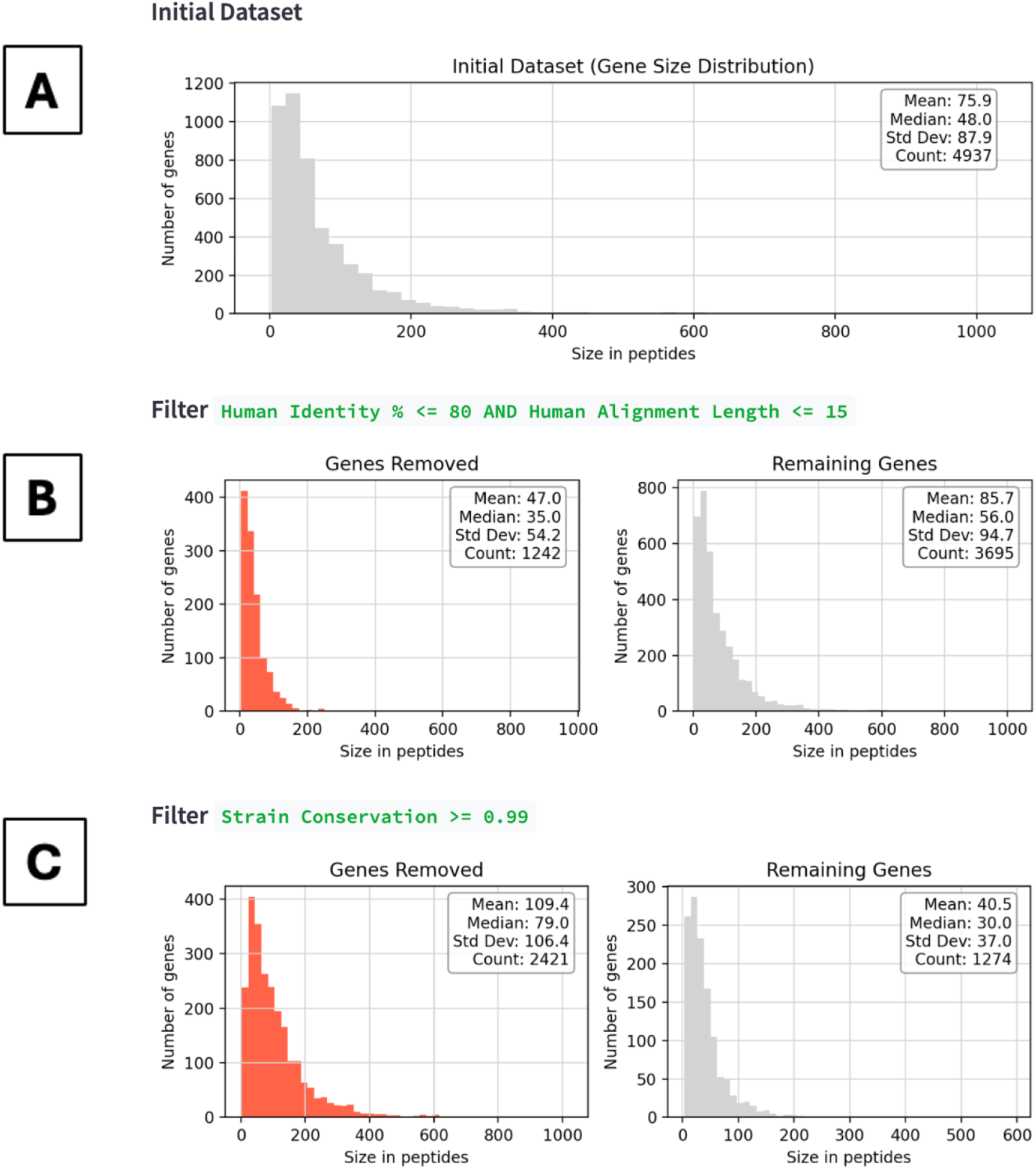
Histograms allow the user to visually explore how chosen filters influence the distribution of gene sizes in the candidate set. Panel A shows the initial gene candidates included in the dataset, while subsequent panels (B and C) illustrate the effect of different filtering options (human identity and strain conservation respectively). In Panels B and C, each histogram displays the size distribution of genes that were removed by a specific filter (left), as well as the size distribution of the remaining genes after each filter has been applied (right). As expected, excluding genes based on strain conservation is more stringent on longer genes, because on average, these have more positions in total which vary. By examining these histograms and associated statistics, these visualisations help the user interpret how filtering decisions shape the gene candidate pool, enabling adjustment of the filters according to requirements.

## Discussion

In this study, we present Pf-PeptideFilter, a customisable web-based tool for the identification and prioritisation of vaccine candidates in *Plasmodium falciparum*. By integrating population-scale genomic data, transcriptomic profiles, and cross-species similarity, Pf-PeptideFilter enables the systematic filtering of potential antigens for vaccine development. Through a modular interface, users can dynamically apply a series of biologically relevant filters–including strain conservation, similarity to human proteins, gene expression, indel frequency, and cross-species homology, either individually or in combination. Using a series of example analyses, we demonstrated how these filters shape resulting candidate sets, enabling the generation of focused shortlists for downstream experimental validation. Pf-PeptideFilter is available as an interactive Streamlit app, providing an accessible and scalable framework for rational vaccine design (https://pf-peptidefilter.streamlit.app/).

By leveraging large genomic datasets to filter potential candidates, Pf-PeptideFilter addresses one of the major obstacles in the development of effective malaria vaccines, that of immune escape through parasite genetic diversity (26,48). Previous vaccine efforts have repeatedly encountered issues with limited efficacy due to antigenic variation among parasites, including trials targeting antigens such as circumsporozoite protein (CSP), apical membrane antigen 1 (AMA1), or merozoite surface protein 1 (MSP1). These vaccines have often resulted in strain-specific immune responses, leading to a lack of broad-population immunity and localised immune escape (27,49). By incorporating genomic variation during the early stages of vaccine selection, Pf-PeptideFilter reduces the likelihood of such outcomes in future. The database and interface also address further challenges in vaccine candidate prioritisation, by reducing autoimmune cross-reactivity risk (via human sequence similarity filtering), ensuring biological relevance (through liver-stage gene expression), and enhancing experimental validation (by prioritising genes with orthologues in model *Plasmodium* species). Together, these improvements provide a systematic, rational approach to vaccine candidate selection for *P. falciparum*.

A key strength of Pf-PeptideFilter is its extensive customisability. Users can precisely control filter stringency using percentile-based thresholds, allowing tailored exploration of potential candidates. In our case study, applying the five core filters at baseline thresholds yielded a shortlist of 253 genes and 10,088 peptides, a set sufficiently narrow for experimental follow-up, yet broad enough to maintain candidate diversity. Visual diagnostics within the app help users interpret how filters affect candidate selection, providing transparency and supporting informed decision-making (21–23), while the inclusion of enrichment summaries for cellular localisation helps users to interpret how filtering decisions shape the composition of candidate sets. Importantly, its user-friendly design ensures accessibility for researchers with minimal bioinformatics experience, facilitating integration into workflows such as peptide library synthesis, functional assays, or T-cell stimulation experiments.

In our example case, the approach recovered several proteins which have previously been shown to have immunogenic properties, such as TCTP and SPATR, the latter of which is a liver-stage antigen relevant to hepatocyte invasion (50,51). Under default settings, Pf-PeptideFilter also identified profilin (PFN), an actin-binding protein essential for parasite motility and host cell invasion, which has previously been investigated as a possible component in a multicomponent vaccine (52). Together these results validate our filtering strategy and demonstrate Pf-PeptideFilter’s utility in identifying high-priority peptide vaccine targets. In addition to known antigens, the method uncovered several novel candidates, each with potential as vaccine candidates. These include GAPM2, a glideosome-associated protein critical for motility and red blood cell invasion during the asexual blood stage, which to our knowledge has not yet been evaluated as a vaccine target, despite its essential role (53). Pf-PeptideFilter also identified intracellular proteins such as TOM40 (a mitochondrial import channel involved in protein transport), USP13 (a ubiquitin-specific protease involved in protein degradation), and HSP110c (a cytosolic heat-shock protein involved in parasite stress responses) (54–56). The latter of which may enhance immunogenicity, as heat-shock proteins are known to be immune-stimulatory and may elicit responses if sufficiently divergent from human homologues (57,58). Each of these proteins have desirable properties for vaccine candidates, including low genetic variability, essential cellular functions, and stage-relevant expression, making them potential targets for further study. While identification of these genes should not be taken as an endorsement of their viability as vaccine candidates (experimental validation would be needed to confirm their immunological relevance), our results highlight the potential of Pf-PeptideFilter as both a discovery and prioritisation platform in identifying novel peptide vaccine candidates.

Future iterations of Pf-PeptideFilter will focus on enhancing predictive accuracy and biological relevance. Planned updates will incorporate additional liver-stage gene expression and proteomic datasets, as well as extending support to newer genomic resources, such as the Pf8 dataset, which encompasses whole genome sequencing data for 32,995 *P. falciparum* samples (47). Support for additional, user-defined peptide lengths (e.g. 8 to 20 amino acids) will also be enabled, allowing more precise epitope selection and alignment with specific vaccine strategies. Additional filtering capabilities–such as epitope motif prediction and post-translational modifications data–will also be included, allowing users to further refine candidate lists. Lastly, we aim to offer a post-filtering ranking module, which will incorporate HLA-peptide binding predictions based on epitope prediction algorithms (59), as well as functional categorisation of identified peptides and genes based on gene ontology. The app will also use these results to allow downstream analysis of HLA binding results, including clustering of peptides based on BLOSUM-based similarity scores or country-specific alleles, improving the interpretation of peptide binding predictions.

In summary, Pf-PeptideFilter offers a flexible, transparent, and reproducible platform for generating shortlists of *P. falciparum* vaccine candidates. By enabling user-defined exploration of diverse biological data, the tool accelerates early-stage vaccine development by streamlining the integration of computational filtering and experimental testing. As the core-framework is not pathogen-specific, Pf-PeptideFilter’s could be adapted for vaccine discovery in other pathogens, including bacterial or viral species, as well as beyond vaccine design, with its modular filtering approach supporting several diverse applications, including the identification of conserved drug targets, or in designing monoclonal antibody therapies. In summary, Pf-PeptideFilter will continue to evolve as part of a suite of *P. falciparum* genomic exploration tools from MalariaGEN (46), which together contribute to global efforts towards malaria eradication.

## Contributions

Conceptualisation - J.A.G, A.J.B., R.P., C.A, Methodology - J.A.G., A.J.B., R.P., C. A., Formal Analysis - J.A.G., A.J.B., C. L., Figures - A.J.B., J.A.G., Investigation - J.A.G., A.J.B., Writing code - J.A.G., A.J.B., Writing Initial Manuscript Draft - A.J.B., Writing, Reviewing and Editing subsequent drafts - A.J.B., J.A.G., C. A., R.P., C,L., Supervision - J.A.G., C. A., R.P., A.J.B., Funding Acquisition - C. A., J.A.G., R.P.. All authors provided input to the manuscript and reviewed the final version.

## Acknowledgments

We would like to acknowledge Matthias Pauthner of the Gates Foundation for useful comments and advice throughout this project. We would also like to thank John Hausmann (Stanford University) and Jake Baum (University of New South Wales) for feedback and comments on the tool. Lastly, we also would like to acknowledge Nina White, Eyyüb Ünlü, and Kate Rowlands for their comments and suggestions on the manuscript, as well as Ben Jackson for useful comments during project development.

## Data Availability Statement

Pf7 population genomic variation data are publicly available from MalariaGEN (Pf7). Liver-stage transcriptomic data are publicly available from Zanghí et al. (2025). Pf-PeptideFilter is publicly accessible at https://pf-peptidefilter.streamlit.app/. All code and data required to reproduce the app is available at: https://github.com/jalmagro/pep-filter.git.

## Declaration of generative AI and AI-assisted technologies in the writing process

The authors declare that they used ChatGPT 4.0 and 5.1 to improve readability, grammar and syntax during writing of the manuscript, but did not use this tool for content generation. The authors reviewed and edited subsequent drafts and take full responsibility for its final contents.

