## Supplementary Information for "Pf-PeptideFilter: An Interactive Catalogue of Peptide Vaccine Candidates for *Plasmodium falciparum*"

**1. Project overview** **2**

1.1. Key filters 2

**2. How to Run the App Locally** **3**

**3. Details on key metrics** **4**

3.1. Strain Conservation 5

3.2. Human Identity Filtering 5

3.3. Indel removal 5

3.4. Homology across Plasmodium species 5

3.5. Gene Expression 5

**4. Computation and extrapolation of filtering criteria** **7**

**5. Usage guides and tutorial** **9**

5.1. Filtering Mechanism 9

5.2. Example - Strain Conservation Filter 9

5.3. Example - Applying Multiple Filters 11

5.3 Downloading Filtering Results 12

**6. Other details** **13**

### 1. Project overview

The Pf-PeptideFilter app is a bioinformatics tool designed to help identify candidate genes and peptides in *Plasmodium falciparum* for vaccine development. It integrates genomic variation, expression data, and similarity to human proteins to systematically rank potential vaccine targets. Users can apply customisable filters and metrics to refine their selection, generating shortlists for downstream applications such as peptide library design or experimental validation.

#### 1.1. Key filters

- **Sequence Conservation Across *P. falciparum* Strains:** Identifies conserved genes and peptides across *P. falciparum* strains to reduce chance of immune escape and ensure efficacy across parasite populations.
- **Similarity to Human Proteins:** Uses BLAST-based filtering to remove peptides with high similarity to human exons, minimising autoimmune risks.
- **Cross-Species Homology:** Ensures vaccine candidates are relevant across *Plasmodium* species, aiding translational studies.
- **Gene Expression Data**: Prioritises genes expressed during critical infection stages (e.g., liver/sporozoite stages).
- **Indel removal:** Filters out genes with high-frequency insertions or deletions, prioritising stable targets less prone to these mutations.

The output includes both gene-level and peptide-level filtered lists, with users able to download summaries and full datasets. Large datasets (>20K peptides) are compressed for efficient access. Pf-PeptideFilter is open-source and freely available at [https://pf-peptidefilter.streamlit.app/](https://pf-peptidefilter.streamlit.app/).

#### 2. How to Run the App Locally

The associated Github repository (https://github.com/jalmagro/pep-filter.git) for Pf-PeptideFilter contains a Streamlit app. Follow the steps below to clone the repository and run the app locally.

##### Prerequisites

Ensure you have the following installed:

- Python 3.7 or higher
- `pip` (Python package installer)
- Dependencies specified in `requirements.txt`:
  - `streamlit==1.35.0`
  - `pandas==1.5.1`
  - `numpy>=1.24.2`

##### 1. Clone the Repository

Open your terminal or command prompt and run:

```
git clone git@github.com:jalmagro/pep-filter.git
cd pep-explorer
```

##### 2. Create and Activate a Virtual Environment (Optional but Recommended)

```
python -m venv .venv
```

- On **Windows**:
  ```
  .venv\Scripts\activate
  ```
- On **macOS/Linux**:
  ```
  source .venv/bin/activate
  ```

##### 3. Install Dependencies

```
pip install -r requirements.txt
```

##### 4. Ensure Data Availability

Ensure the datasets:

`gene-metrics-filtering.csv.gz` and `peptide-metrics-filtering.csv.gz` are located in the `data` directory within the project.

##### 5. Run the App

Start the Streamlit app using the following command:

```
streamlit run app.py
```

The app will launch locally and open in your browser.

##### 6. Access the App

Or, open your browser and navigate to:

`http://localhost:8501/ or https://pf-peptidefilter.streamlit.app/`

##### Contributing

We strongly encourage feature requests! Please open an issue on this repository or submit your requests via our Google Forms.

##### Licence

This project is licensed under the ‘MIT License’. See the LICENCE file for more details.

### 3. Details on key metrics

#### 3.1. Strain Conservation

Pf-PeptideFilter assesses conservation at both the gene and peptide levels, using MalariaGEN's Pf7 dataset (~16,000 genomes). Conservation is calculated using:

- The frequency of identical amino acid haplotypes across ~8,500 African samples.
- Sequence identity compared to the 3D7 reference genome.

Higher conservation thresholds reduce the risk of immune escape and ensure efficacy across diverse populations

#### 3.2. Human Identity Filtering

To prevent cross-reactivity, peptides are BLASTed against human exons, estimating both percentage identity and alignment length. For example, a 60% identity score corresponds to an 80% match over an alignment covering 75% of the peptide. Genes are scored based on the mean identity of their constituent peptides, excluding genes or peptides that closely resemble human proteins.

#### 3.3. Indel removal

Genes with high-frequency insertions or deletions (especially frameshifts) are excluded to prioritise stable targets, as these could compromise vaccine efficacy. Indel frequencies are calculated across Pf7 samples.

#### 3.4. Homology across *Plasmodium* species

Genes without orthologs in *P. vivax, P. berghei*, or *P. knowlesi* can be excluded. Genes are filtered based on one-to-one ortholog correspondence between *P. falciparum* and these species. This feature can enable selection of targets relevant to animal models.

#### 3.5. Gene Expression

Gene expression filtering prioritises genes actively expressed during critical infection stages, such as the liver and sporozoite stages. Using data from Zanghi et al. (2025), genes pass if all replicates have CPM ≥1 in liver-stage samples (days 2, 4, 5, and 6). Peptides are included only if their associated gene passes the filter. This ensures that identified targets are accessible to the immune system during key points of infection, improving their relevance for vaccine development.

### 4. Computation and extrapolation of filtering criteria

Filters in Pf-PeptideFilter are applied independently, either at the gene or peptide level, depending on the metric. This avoids biases that could be introduced by aggregating gene-level data by only assessing peptides. In some cases, metrics computed for peptides are extrapolated to their corresponding genes, enabling flexibility in filtering based on research priorities (Supplementary Table 1).

**Supplementary Table 1.** Detail on computation and extrapolation of filtering metrics used in Pf-PeptideFilter.

| **Category** | **Criterion** | **Level at which computed** | **Extrapolation for genes** | **Extrapolation for peptides** | **Remarks** |
|---|---|---|---|---|---|
| Filtering | Strain Conservation | Both peptide and gene levels | None | None | Full exons are longer than peptides (20-AAs), making them more likely to have more mutations and stratify into more haplotypes. Using genes rather than peptide filtering therefore has the effect of reducing the relative frequency of the 3D7 haplotype.<br><br>**Optional:** assign to a gene the average 3D7 frequency of all its peptides. |
| | Human Identity (BLAST) | Peptide (against human exons) | Averaged metric for all peptides in the gene | None | Averaging dilutes the signal, as most peptides do not have matches in human exons. However, computing matches at gene level would not work as they do not have matches.<br><br>**Optional:** assign to each gene the maximum metric value of any of its peptides (i.e. the peptide with the best match. |
| | Indel removal | Both peptide | None | None | Targets with high indel |

| | | | | | |
|---|---|---|---|---|---|
| | | and gene levels | | | frequencies, particularly frameshifts, are excluded to avoid rapidly mutating candidates. |
| | Homology | Gene | None | Ortholog if the gene it belongs to is an ortholog | Gene-level filtering only. One-to-one ortholog correspondence ensures cross-species applicability. |
| | Gene Expression | Gene | None | Expressed if the gene it belongs to is expressed | A peptide passes the filter only if its associated gene is actively expressed. |

### 5. Usage guides and tutorial

We generated a custom database of information on genetic variation in *P. falciparum*, built upon the publicly accessible Pf7 dataset of whole genome sequences, and aggregated by MalariaGEN. There are 14 chromosomes in the *P. falciparum* genome, consisting of a total of 4,937 genes. Calculating each 20 amino acid peptide using a sliding window (sliding every 10 amino acids), gives a total of 379,325 peptides across the genome. The app can be used to apply a series of filtering criteria to this database, allowing the user to filter candidate genes/peptides based on their preferred combination of criteria–such as those which could be used as potential vaccine targets.

#### 5.1. Filtering Mechanism

The database is built at the percentile level, meaning that for each numeric filter, users can specify the minimum percentage of gene peptides that need to pass the filter. The app supports threshold values ranging from 50% to 100%, along with several intermediate options (60%, 70%, 75%, 78%, 80%, 82%, 85%, 88%, and whole numbers between 90-100%). This allows for flexible and precise filtering tailored to different research objectives.

#### 5.2. Example - Strain Conservation Filter

Below is an example using the strain conservation filter. When using the filter at a minimum frequency of 0.99, the number of retained genes will depend on what percentage of the gene's peptides meet this threshold. If 95% of the peptides must pass the test at a 0.99 frequency, the app will retain just over 2,000 genes. Notice there are fewer genes retained if we increase the stringency of the filter (Supplementary Figure S1).

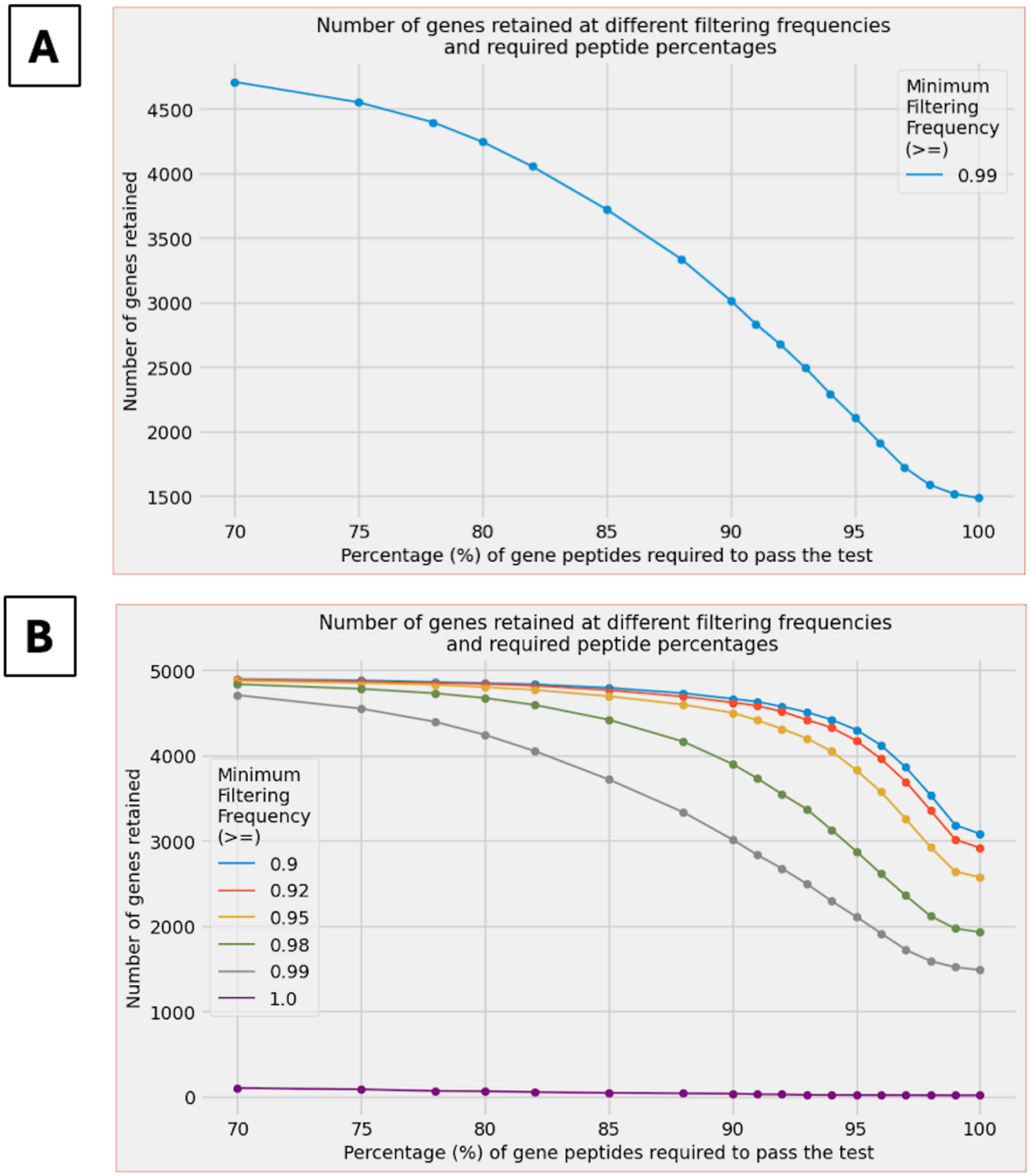


**Supplementary Figure S1.** A) Decay curve illustrating the impact of the strain conservation filter at different stringencies. Note, as the percentage of gene peptides required to pass the test increases, the number of genes retained by the filter decreases substantially, even though the minimum filtering frequency remains constant. B) Increasing the required percentage of peptides passing the strain conservation test will result in a gradual reduction of retained genes, with the curve becoming steeper above the 90% threshold. For example, the decay curves are steeper at higher frequencies.

#### 5.3. Example - Applying Multiple Filters

The example provided so far only uses a single filtering metric. Applying filtering using two separate criteria gives a smaller list of candidates (Supplementary Figure S3):

- Human identity filtering (95% of peptides in a given gene need to have less than 60% identity over an up to 20 amino acid alignment)
- Strain conservation (95% of peptides in a given gene need to be representative of the dominant haplotype (0.99), irrespective of 3D7).

| filter | remaining | fraction |
|---|---|---|
| Initial Dataset | 4,937 | 1 |
| Human Identity % <= 60 AND Human Alignment Lenght <= 20 | 2,171 | 0.4397 |
| Strain Conservation >= 0.99 | 804 | 0.1629 |

**Human identity gene-extrapolation rule:** `Minimum percentage of gene peptides`

- Up to **5% of gene peptides can fail** identity and length filters

**Strain conservation gene-extrapolation rule:** `Minimum percentage of gene peptides`

- At least **95% of gene peptides must pass** frequency filter

**Genes retained:** `804`

**Total number of peptides:** `28,204`

**Average number of peptides per gene:** `35.08`

**Supplementary Figure S3.** Effect of simultaneously filtering on two independent criteria (strain conservation and human identity). Applying multiple filters drastically reduces the number of retained genes. These candidates can then be exported as lists, in the form of an Excel or CSV file.

#### 5.3 Downloading Filtering Results

Once filtering criteria have been chosen, the user then has the option to download the summary of the applied filters and the final list of filtered genes (for results with more than 20K peptides, the output file is compressed with gzip). When downloading the gene data, the following columns relate to AA haplotypes:

- gene_haplotype - the most frequent AA haplotype found for the whole gene.
- peptide_list - the list of most frequent AA haplotypes found for each gene peptide (separated by commas).

Note that genes and peptides are filtered independently (at the gene and peptide level) but some metrics are extrapolated (see above). The peptide_list includes all the peptides in the gene (not only those that pass all filters if you were filtering by percentage of gene peptides) and these can be different to the sequences in gene_haplotype**.**

#### 6. Other details

**Using Mean Values Over Gene Peptides:** This method calculates the average of the metrics across all peptides in a gene. While this approach may dilute the signal (since many peptides may not have matches), it does provide an overall metric for the gene.

**Using Percentile Values Over Gene Peptides:** Alternatively, the user can represent a gene using a specific percentile value calculated across all its peptides. This allows for more granular control over how the gene is represented (Supplementary Table S2).

**Supplementary Table S2.** Examples of percentile thresholds for use in filtering metrics.

| Percentile Examples | Gene Representation |
|---|---|
| **0th Percentile** | The smallest observed metric value across all peptides. |
| **50th Percentile** | The median metric value. |
| **100th Percentile** | The largest observed metric value. |
| **Xth Percentile** | The metric value for which X% of peptides have values ≤ X (e.g., the 90th percentile includes 90% of the peptides). |

For example, if a user sets the alignment length slider to 18 amino acids (AAs) and applies the alignment length filter at the 90th percentile, the peptides of each gene will be sorted by alignment length in increasing order. The filter will then select the length that is greater than 90% of the peptides for each gene. This value will therefore represent the gene in the filter evaluation. As a result, any gene for which 10% or more of its peptides have a matching alignment length exceeding 18 AAs (the threshold set by the slider) will be removed.

The same applies to strain conservation, the user can choose between using a single full-gene haplotype representing the gene for frequency filtering (previous behaviour) or to use a percentile frequency from its peptides. For example, if the user sets the minimum allowed frequency to 0.99 and sets frequency percentiles to the $10^{th}$ percentile, they will filter out any gene for which less than 90% of their peptides have a frequency below 0.99. This kind of filtering, which appears convoluted at first, allows for finer control on how to filter genes from their peptides. The rationale is that there are many genes for which the peptide metrics statistics could be very heterogeneous. This allows the user to specify what fraction of peptides in a gene they are willing to tolerate failing the filter without losing the gene. This is statistically more robust than blindly relying on the mean.
